# Variation in the strength of selection but no trait divergence between elevational extremes in a tropical rainforest *Drosophila*

**DOI:** 10.1101/2021.08.15.456429

**Authors:** Eleanor K. O’Brien, Megan Higgie, Andrew D. Saxon, Ary A. Hoffmann, Jon Bridle

**Affiliations:** School of Biological Sciences, Life Sciences Building, University of Bristol, Bristol BS8 1TQ, United Kingdom; Centre for Precision Health, School of Medical & Health Science, Edith Cowan University, WA 6027, Australia; College of Marine & Environmental Sciences, James Cook University, Queensland 4811, Australia; Bio21 Institute, University of Melbourne, Victoria 3052, Australia; Department of Genetics, Environment & Evolution, University College London, London, United Kingdom

**Author notes:** **Author email addresses:** EKO, MH, ADS, AAH, JRB. **Corresponding author:** Eleanor K. O’Brien, Centre for Precision Health, School of Medical & Health Science, Edith Cowan University, WA 6027, Australia.

**Keywords:** caged transplant experiments, *Drosophila*, elevation gradient, fitness, genetic variance, heritability, quantitative traits, selection

## Abstract

Evolutionary responses to environmental change require heritable variation in traits under selection. Both heritability and selection vary with the environment, and may also covary, so environmental variation can affect evolutionary rates. However, simultaneous estimates of heritability and selection across environmental gradients in the field are rare. We estimated environmental variation in selection on three traits (cold tolerance, heat tolerance, and wing size) of the rainforest fruitfly *Drosophila birchii*. We transplanted flies in cages along two elevational gradients in north-east Queensland, Australia, and calculated the genetic covariance of trait values with fitness (productivity in cages) at each site. For wing size, we estimated heritability at each site from the correlation between field-reared offspring and their laboratory-reared relatives. We used estimates of selection and heritability to predict selection responses along the elevation gradients, and compared these with trait variation observed in flies sourced from populations at elevational extremes. We found evidence for divergent selection on cold tolerance across elevation at one gradient. Heritability of wing size was highest at gradient ends, and the size of the predicted response to selection on this trait increased with elevation. However, the direction of selection varied, even among adjacent sites, making such selection unlikely to cause divergence of this trait with elevation. None of the traits differed between populations from elevational extremes at either gradient. Variation in the strength and direction of selection over small spatial scales, as well as in time, may explain why predicted responses to selection are often not seen in natural populations.

## Introduction

Climate change and habitat loss are altering abiotic and biotic environmental conditions and causing global biodiversity declines (Sala et al., 2000, Urban, 2015, Nadeau et al., 2017). Evolutionary responses to these environmental changes are likely to be crucial for the persistence of many species across their ranges, particularly where populations are unable to migrate to track suitable habitat due to geographic or biological barriers to dispersal (Bridle & Vines, 2007, Hoffmann & Sgrò, 2011). Such adaptive responses require heritable variation in trait(s) under selection, as described by the breeder’s equation (Lush, 1937, Falconer & Mackay, 1996) and its multivariate equivalent (Lande, 1979). However, it remains unclear how environmental change affects evolutionary potential alongside causing spatially and temporally varying selection.

Quantitative genetic analyses of natural populations often reveal significant heritable variation in a diverse range of traits (e.g. Mousseau & Roff, 1987, Mousseau et al., 2000, Kruuk et al., 2008), and provide examples of apparently strong selection on many traits (Kingsolver et al., 2001, e.g. Kruuk et al., 2002, Hereford et al., 2004), suggesting that the potential for adaptive evolution can often be high. However, evolutionary responses in nature are often weaker or even in the opposite direction to those predicted (Kruuk et al., 2002, Postma et al., 2007, Wilson et al., 2007, e.g. Gienapp et al., 2008, Pemberton, 2010). Explanations for this include correlations with other (unmeasured) traits (Blows & Hoffmann, 2005, Walsh & Blows, 2009), effects of spatial and temporal environmental variation in the field on both heritability (Hoffmann & Merilä, 1999, Charmantier & Garant, 2005) and the strength of selection (Siepielski et al., 2013, Siepielski et al., 2017, Hayward et al., 2018), and low levels of heritable variation in traits that are ecologically critical (Blows & Hoffmann, 2005). These issues mean that estimates of heritability and selection made under constant conditions in the laboratory will often be of little use in predicting evolutionary responses in the field.

Direct effects of the environment on traits (e.g., through condition dependence, or plasticity, or both) can affect their heritabilities, although it is unclear when heritability will be increased or decreased in less favourable environments (Hoffmann & Merilä, 1999, Charmantier & Garant, 2005, Rowiński & Rogell, 2017). Selection on a trait or combination of traits will also change between environments due to changes in the relationship of the trait(s) with fitness (Siepielski et al., 2013, Siepielski et al., 2017, Hayward et al., 2018). In addition, the environment itself may create a correlation between heritability and selection, which could constrain or increase the rate of evolution along particular trajectories (Wilson et al., 2006, Husby et al., 2011, Wood & Brodie III, 2016, although see Ramakers et al., 2018). For example, in Soay sheep (*Ovis aries*), heritability and selection on birthweight were negatively correlated due to dependence of both parameters on changes in the environment among years (Wilson et al., 2006). Specifically, heritability increased but the strength of selection decreased in years when conditions were more favourable (i.e. years when lamb survival was higher). By contrast, when selection for high birthweight was stronger (i.e. when lamb survival was lower), heritability of birthweight was reduced. Adaptive evolution of birthweight is therefore constrained by weaker selection in good years and by lower heritability in harsh years (Wilson et al., 2006). By contrast, Husby et al. (2011) found a positive correlation between heritability and selection on the timing of breeding in great tits (*Parus major*), which may increase the evolutionary potential of this trait to ongoing climate change.

Estimates of changes in heritability and selection along environmental gradients in the field are scarce in animals, and are almost entirely restricted to vertebrates (Wilson et al., 2006, Quinn et al., 2009, Husby et al., 2011, Hayward et al., 2018). Field transplant experiments are a powerful method to test the effect of spatial variation in the environment on evolutionary potential, by enabling selection differentials and heritabilities of traits to be estimated on the same set of genotypes or families at different sites along an environmental gradient (e.g. Etterson, 2004). O’Brien et al. (2017) used a caged transplant experiment to estimate genetic and environmental variation in fitness of the tropical rainforest specialist *Drosophila birchii* along elevation gradients in north Queensland, Australia. Elevation is strongly associated with climate variables, including mean, minimum and maximum temperature and humidity, and with the abundance and fitness of *D. birchii,* making transplants along this gradient an ecologically relevant test for existing local adaptation in the face of climatic change. These data showed that fitness of *D. birchii* in cages (measured as the number of offspring produced per cage) declined with increasing elevation, despite the field density of *D. birchii* being higher at high than low elevation (O’Brien et al., 2017). This suggests that when the effects of climate are considered without any associated effects on food availability or biotic interactions, high elevation habitats are less productive than low elevation habitats, and that selection varies in strength or direction with elevation for traits such as thermal responses. This study also detected substantial genetic variation in fitness in field cages, and genetic divergence in productivity when isofemale lines sourced from elevational extremes were raised under laboratory conditions, but no evidence for local adaptation to elevation, even when comparing flies that had been reciprocally transplanted between the high and low elevation limits of the species’ elevational range at those latitudes (O’Brien et al., 2017). This system provides an opportunity to test whether evolutionary potential varies across this species’ range due to environmental variation in heritability or selection (or both), which will enable predictions about where and when adaptive responses (e.g. to climate change) are most likely.

In this paper, we estimate selection on, and heritability of, ecologically relevant traits of *D. birchii* at sites along two elevation gradients that span the elevational range of this species. We assayed relatives of the flies transplanted in field cages in O’Brien et al. (2017) to obtain family means for three traits potentially under selection along these elevation gradients: cold tolerance, heat tolerance and wing size. We estimated the genetic (co)variances within and between these traits and used the covariance of trait means (measured in the laboratory) with fitness of the same lines in cages in the field to estimate selection on each trait at each elevation. For wing size, we also tested how heritability varies with environment, using a regression of the wing size of daughters emerging from field cages at each elevation on that of their laboratory-reared relatives, and applied the breeders’ equation to predict the direction and magnitude of the response to selection on this trait at sites along each elevation gradient. We then compared our predicted responses with trait divergence between populations from elevational extremes to determine whether the responses to selection that should occur based on trait covariance with fitness are realised in the field.

## Methods

### Collection and establishment of *D. birchii* isofemale lines for quantitative trait assays and field transplant experiments

We established isofemale lines (hereafter ‘lines’) of *Drosophila birchii* from field-mated females collected from banana baits in rainforest at sites along two elevation gradients: Mt Edith (17° 7.9’S, 145°37.2’E) and Paluma (18°59.0’S, 146° 14.0’E) in north Queensland, Australia, using the same methods described in Bridle et al. (2009) and O’Brien et al. (2017). We established five *D. birchii* lines from each of eight populations: two high and two low elevation populations from each of the two gradients. These gradients are characterised by a warmer, drier, more fluctuating environment at low elevations, and a cooler, wetter, more stable environment at high elevations (see O’Brien et al., 2017 and Table 1 for temperature and humidity data at each gradient). Our sampling represents the full elevational range of this species at these latitudes (0-1100 m above sea level), therefore it should capture the full range of phenotypic and genetic diversity in traits associated with fitness under these different climatic conditions. The lines from each population were crossed with one another (see Appendix S1 and O’Brien et al. (2017)) to generate flies for use in field caged transplants, and for estimates of thermal tolerances and body size. Lines maintained in the laboratory were held at 25 °C on a 12:12 hr light:dark cycle unless otherwise stated.

**Table 1.** Location (latitude and longitude), elevation (in m above sea level), length, number of transplant sites and mean temperature and humidity at the two elevation gradients used in the caged transplant experiment and as the source of *D. birchii* for field and lab assays.

| Gradient | Latitude (°S) | Longitude (°E) | Elevation range (m) | Total length (km) | No. of transplant sites | Mean daily temperature range (°C) | Mean daily relative humidity range (%) |
| --- | --- | --- | --- | --- | --- | --- | --- |
| Mt Edith | 17°7.9' | 145°37.2' | 697–1105 | 4.3 | 9 | 13.6–15.2 | 95.4–99.5 |
| Paluma | 18°59.0' | 146°14.0' | 72–916 | 3.7 | 10 | 14.7–19.7 | 74.3–78.3 |

### Field cage transplant experiment

In May 2012, 5910 flies emerging from the line crosses were transplanted in 591 cages to 19 sites along the two elevational gradients from which they were collected (325 cages across 9 sites at Mt Edith, and 266 cages across 10 sites at Paluma; Table 1). The details of the experimental design are described in O’Brien et al. (2017) and summarised briefly below.

The transplant sites spanned the full elevation range of this species and included the low and high elevation sites from where lines originated. Because it was not feasible to replicate every line-cross combination at every site, offspring of crosses from the same maternal isofemale line were combined prior to cage establishment by taking equal numbers of offspring from each cross. In total, we transplanted 15 lines at Mt Edith and 20 lines at Paluma. Lines were only transplanted back into their gradient of origin, not between gradients. Each cage contained five virgin males and five virgin females (all aged 3–10 days) from the same maternal line, and each maternal line was represented by 2–4 cages per site. Adult flies were left in cages to mate and lay for five days and then removed. Cages were left in the field for an additional 25 days (30 days total) to ensure all offspring had emerged at all sites. Because flies were placed in cages as virgins, all courtship, mating, egg laying and offspring development took place under field conditions. The number of offspring emerging from each cage was recorded and used as our measure of fitness. Analyses of variation in cage fitness (productivity) along these gradients have been published previously (O’Brien et al., 2017). Here, we assayed cold tolerance, heat tolerance and wing size on descendants of flies transplanted in field cages, and we used productivity in cages to estimate selection on these traits at each transplant site.

### Laboratory assays of quantitative traits

Using flies emerging from crosses between the same *D. birchii* lines that were transplanted into cages in the field experiment, we assayed three traits: cold tolerance, heat tolerance and wing size. Similar traits show repeated genetic divergence along thermal (latitudinal and elevational) gradients in several *Drosophila* species (Hallas et al., 2002, e.g. Hoffmann et al., 2002, Sørensen et al., 2005), including cold tolerance in *D. birchii* (Bridle et al., 2009). For wing size, we used flies from the same set of crosses (and generation) used to generate flies for the transplant experiment. Cold tolerance and heat tolerance were assayed on flies emerging from crosses between the same lines, three generations after the transplant experiment.

Cold tolerance was measured as the productivity (number of offspring emerging from three days of laying) of six-day old virgin females subject to a cold shock that induced a chill coma, then mated to a single male; heat tolerance was measured on seven-day old virgin females as the time to lose standing ability on exposure to a static heat shock of 37 °C; wing size was measured as the centroid size of the right wing of female flies that were at least two days of age, calculated from the positions of 10 landmarks (Figure S1; Griffiths et al. (2005)). See Appendix S2 for a detailed description of trait assays.

#### Data analysis

All analyses were run using R v. 4.1.0 (R Core Team, 2020), executed in R Studio v. 1.4.1103 (RStudio Team, 2021). Details of specific packages and functions used are given in the description of each analysis.

### Divergence of traits within and between elevation gradients

We tested for divergence of traits (cold tolerance, heat tolerance and wing size) between *D. birchii* sourced from the two elevation gradients (Mt Edith and Paluma) and between high and low elevation source populations within each gradient, by fitting linear mixed models using *lme4* v. 1.1-23 (Bates et al., 2015). Separate models were fitted for each trait. Gradient, elevation and their interaction were all fitted as fixed effects. Rearing vial was fitted as a random effect for all traits and batch was additionally included as a random effect in models of heat tolerance. The significance of fixed effects was determined using *F*-tests, and of random effects using a χ^2^ test to compare the log likelihood of models with and without the factor included.

### Estimating additive genetic (co)variances of traits from laboratory crosses

We estimated additive genetic (co)variances of the three traits in the laboratory crosses using an animal model approach (Wilson et al., 2010, Kruuk, 2004), implemented using MCMCglmm (v. 2.24) (Hadfield, 2010). We fitted the following linear model for each gradient:

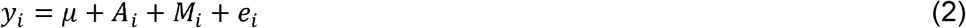

where *y_i_* is the multivariate phenotype of individual *i, μ* is the multivariate phenotypic mean, *A_i_* is the additive genetic effect (fitted as a random effect), *M_i_* is the maternal effect variance (variance among mothers, beyond that explained by additive effects, fitted as a random effect) and *e_i_*, is the residual (error) variance. Values of each trait were standardised within each gradient to a mean of 0 and a standard deviation of 1. Standardisation was performed because large differences between traits in their means and variances can bias estimates of covariance (Careau et al., 2015, Hansen & Houle, 2008). Additive genetic effects were fitted as a matrix of predicted additive covariances between all individuals (i.e., a matrix of the pairwise coefficients of relatedness between individuals). Relatedness was estimated from a pedigree constructed based on our breeding design. See Appendix S3 for a full description of the assumptions made in pedigree construction. Our data set was not large enough to estimate additive genetic (co)variances separately for each source elevation or population, although we did initially run models with source population included as a fixed factor, to account for differences in trait means between populations. However, this had no effect on (co)variance estimates, and so we present results from the simpler models run without this parameter.

We used weakly informative inverse-Wishart priors, where the scale parameter *V* was a 3 x 3 matrix, with phenotypic variance in each trait equally distributed between additive genetic, maternal effect, and residual variation, and the scale parameter *nu* equal to the number of traits. We constrained the residual covariances among traits to be 0, since traits were measured on different individuals. We ran models for 2.5 x 10^6^ iterations, with a burn-in of 5 x 10^5^ iterations and a thinning interval of 2000, giving a total sample size of 1000. This resulted in autocorrelation of successive runs below 0.1 and effective sample sizes close to the maximum for all parameter estimates. Model performance was also evaluated by examining plots of the time series of the values obtained for the posterior probability distribution, and the density of this distribution for all parameters estimated from each model.

We obtained the additive genetic (co)variance of each trait and pairwise trait combinations from the mode of the posterior probability distribution for the relevant model term. We calculated heritability (*h*^2^) as the additive genetic variance (*V*_A_) as a proportion of total phenotypic variance (*V*_P_). We converted genetic covariances to correlations using the *posterior.cor* function in MCMCglmm. We obtained the Highest Posterior Density (HPD) interval around each parameter estimate, and values derived from model parameter estimates. This defines the bounds of the region of the posterior distribution where 95% of samples lie. Additive genetic covariances and correlations between traits were deemed significant if the HPD interval did not overlap zero. Variances are constrained to be above zero, meaning that this method cannot test the significance of additive genetic variances, however we can test whether they differ significantly between gradients or traits by testing for overlap of the HPD intervals.

### Estimating selection on thermal tolerance and wing size of *D. birchii* along elevation gradients

We estimated linear selection differentials (*S*) for cold tolerance, heat tolerance and wing size from a regression of standardised (mean = 0, SD = 1) trait values for each line estimated from laboratory assays against relative fitness in cages at each site. Separate models were fitted for each transplant site, as well as overall models for each gradient (using fitness at all transplant sites), which additionally included transplant site as a random factor. Using standardised phenotypes rather than the raw data means that *S* can be compared across traits and between sites and gradients (Kingsolver et al., 2001).

Fitness was measured as the number of offspring emerging from each cage (see O’Brien et al., 2017), and was converted to relative fitness by dividing this by the mean number of offspring produced in all cages at that transplant site (or gradient in the case of the overall models for each gradient) (Linnen & Hoekstra, 2009). *S* therefore gives the change in relative fitness expected with an increase of one standard deviation in the value of a given trait.

We tested for a linear relationship between selection and elevation (which is expected to drive trait divergence between high and low elevations), by running linear models regressing estimated selection differentials on elevation for each trait at each gradient using the *lm* function in R.

### Variation in heritability of wing size along elevation gradients

We obtained site-specific estimates of heritability of wing size along the two elevation gradients using regressions of the mean wing size of female offspring emerging from field cages on that of females from the same maternal isofemale line emerging from laboratory crosses (i.e., the same pool of flies transplanted as adults in cages). Note that heritability estimated using this method includes maternal genetic (and environmental) effects, therefore we treat it as an overall measure of heritable variation and denote it *H*^2^. Regressions were performed within each transplant site at Mt Edith and Paluma, using the *lm* function in R.

Mean wing size of the female offspring developing in cages was 7.5% larger than females from the laboratory crosses, and the wing size of flies emerging from cages declined with increasing elevation (Figure S2). To prevent differences in trait means between generations and sites from skewing heritability estimates (Careau et al., 2015, Hansen & Houle, 2008), we first standardised wing size data within each group (lab-reared and field-reared) to a mean of 0 and a standard deviation of 1 within each transplant site.

To test for a linear association between heritability and elevation, we ran linear models that regressed field heritability of wing size against elevation at each gradient. We also regressed field heritability of wing size against the selection differential (*S*) for this trait within each gradient, to test for covariation of heritability and the strength of selection. Both of these regression analyses were conducted using the *lm* function in R.

### Estimating response to selection on wing size along elevation gradients

To estimate the response (*R*) to selection at each elevation, we applied a broad interpretation of the breeders’ equation:

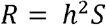

where *S* is the site-specific estimate of the selection differential, and *h*^2^ is the estimated heritability of the trait. In our case we do not have an estimate of *h*^2^ but instead use *H*^2^, acknowledging that this includes maternal effects and other nonadditive variation. We limited this analysis to wing size because this was the only trait for which we had site-specific estimates of *H*^2^. We calculated *R* at each elevation using the estimates of *H*^2^ from the regressions of wing size of lab-reared females and their field-reared female relatives in cages at each site. This allowed us to evaluate the effect of environmental variation in *H*^2^ (as well as in *S*) on responses to selection along elevation gradients for this trait. It also enabled us to compare estimates of *R* that used site-specific estimates of *H*^2^ versus those using a single laboratory estimate. We used the standard errors of *S* (substituted into the breeders’ equation) to estimate standard errors around the selection response *R*.

## Results

### Genetic divergence of thermal tolerance and wing size between, but not within, elevation gradients

*D. birchii* lines from the colder, higher elevation Mt Edith gradient had significantly higher cold tolerance (producing 25% more offspring following a cold-induced coma) and larger wing size (2.2% larger wing size) than those from the warmer, lower elevation Paluma gradient (Figure 1, Table 2) when reared in the laboratory, but there was no significant difference in heat tolerance. None of the three traits differed significantly between low vs. high elevations within either gradient (Figure 1, Table 2).

**Figure 1.**
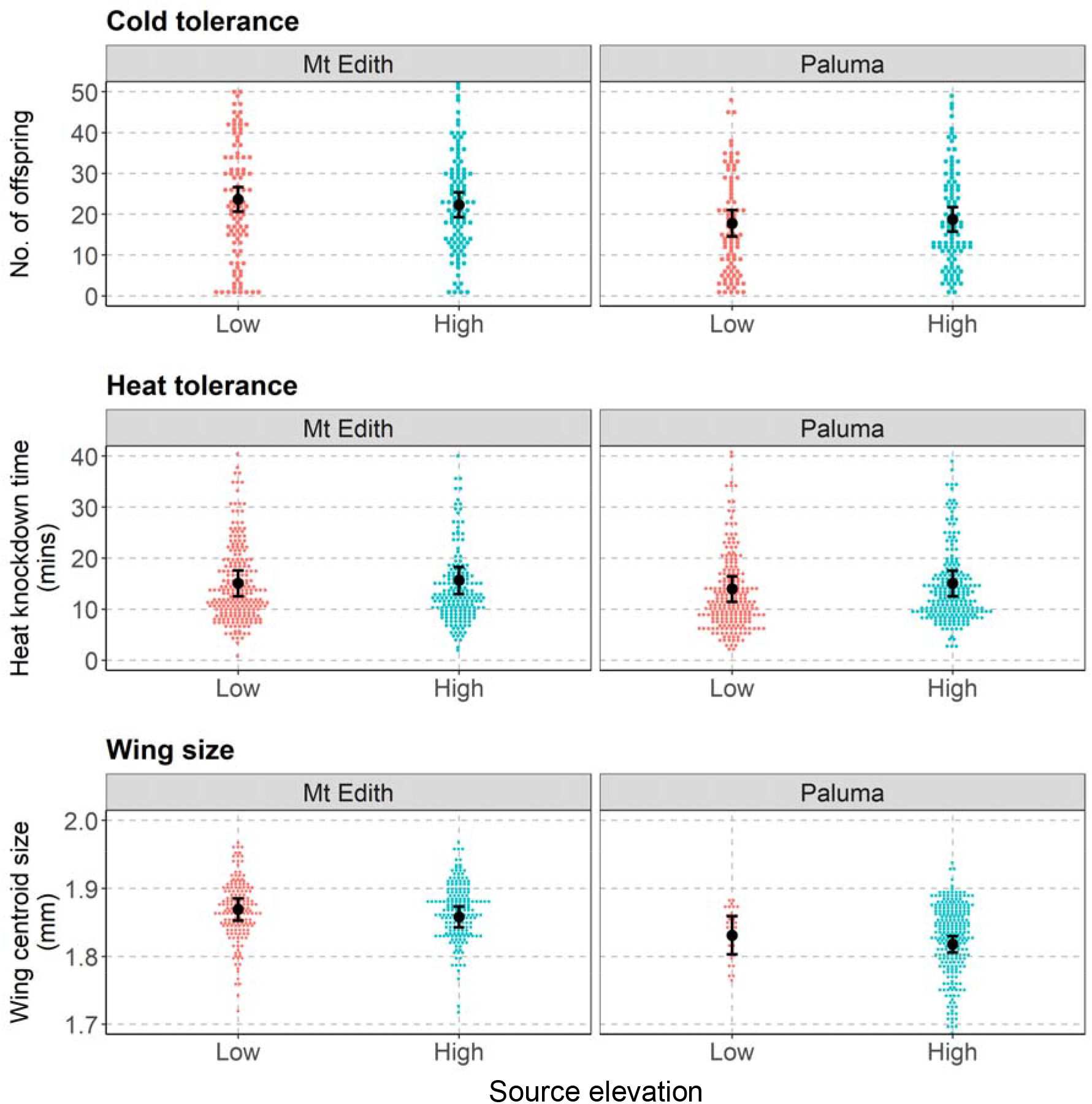
Dot plots showing trait values of *D. birchii* from high (blue dots) and low (red dots) elevation populations at the Mt Edith and Paluma gradients for cold tolerance, heat tolerance, and wing size. Dots are trait values for each individual, plotted in bins equal to ~1/100^th^ of the total range of the data. Dots within the same bin are plotted alongside each other, so the horizontal spread of dots is proportional to the number of individuals within each bin. Black points show estimated marginal means of each trait for each source elevation and gradient from the linear models, with error bars showing the 95% confidence intervals of these estimates. *Drosophila birchii* from the Mt Edith gradient had significantly higher cold tolerance and larger wing size than those from the Paluma gradient. However, we did not detect any difference for heat tolerance, nor any divergence between high and low elevation populations within gradients for any of the three traits (see Table 2).

**Table 2.**
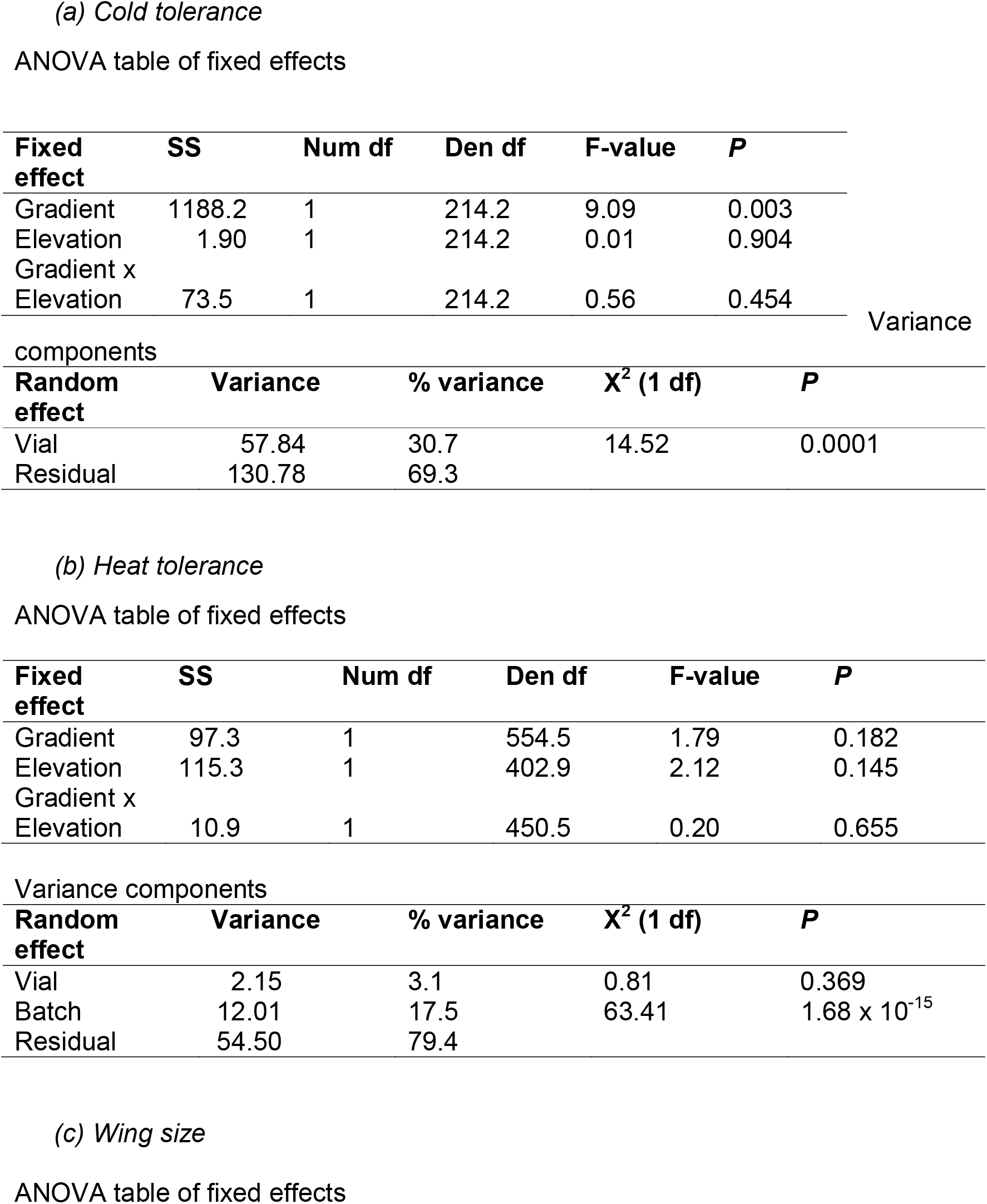
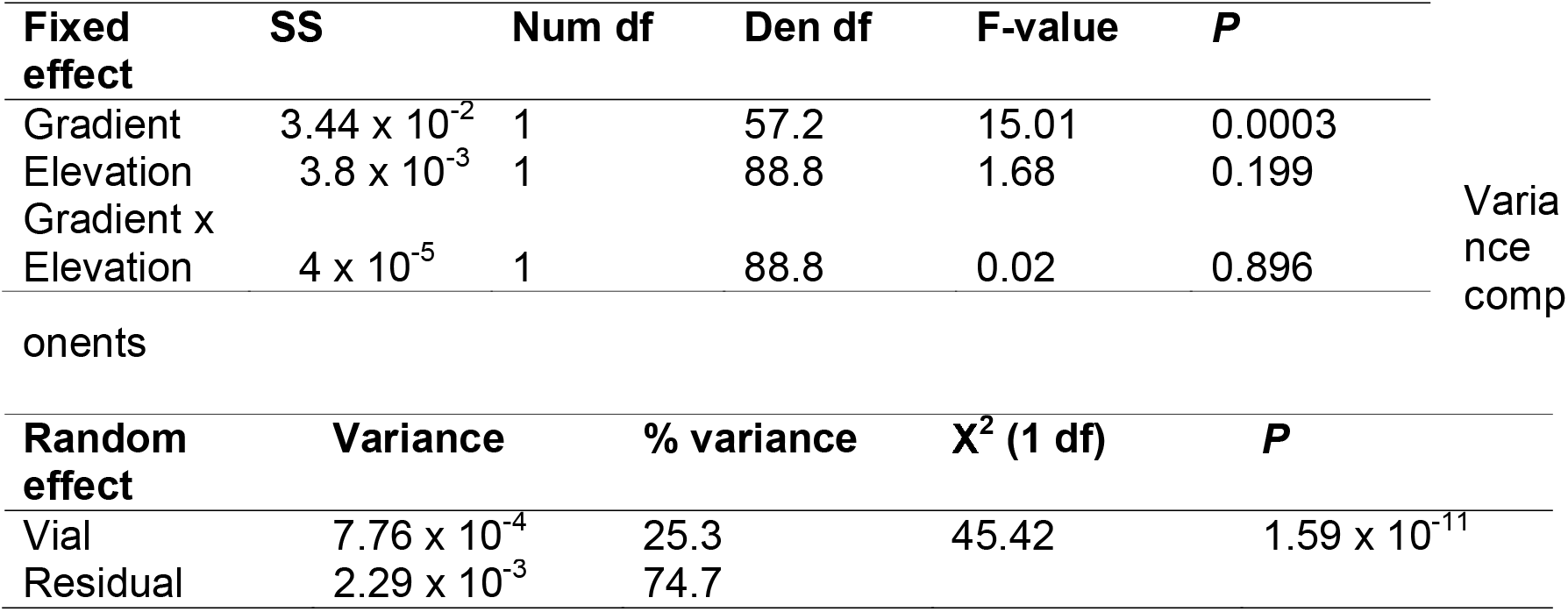
Results of linear mixed models testing for variation in (a) cold tolerance, (b) heat tolerance, and (c) wing size of *Drosophila birchii* sourced from high and low elevation populations at the Mt Edith and Paluma gradients. For each trait, ANOVA tables report the fixed effects of source gradient (Mt Edith vs. Paluma), elevation, and their interaction. Variance component tables show the percentage variance in each trait explained by the random factors in the models (vial and, for heat tolerance, batch), and χ^2^ and *P*-value obtained from a comparison of the log-likelihood of models with and without the factor included.

### Additive genetic (co)variances of thermal tolerance and body size from the laboratory crossing design

Heritabilities of all traits estimated from lab-reared flies emerging from the crossing design ranged from 0.12–0.29 (Table S1). These heritability estimates did not differ between gradients (i.e., the 95% HPD intervals overlap). Maternal effect variance ranged from 0.09–0.22, and also did not differ between gradients (Table S1). At both gradients, estimates of both heritability and maternal effect variance were higher for wing size than for cold or heat tolerance (Table S1) although the 95% HPD intervals overlapped between traits in all cases, so the importance of this difference was unclear.

We did not detect significant additive genetic correlations (*r*_G_) between any of the pairwise trait combinations at either gradient (HPD intervals all overlapped with zero; Table S1). However, these intervals were all extremely wide. Large sampling variance is common when estimating genetic covariances and correlations, meaning very large sample sizes are typically required where trait correlations are weak (Visscher, 1998, Bijma & Bastiaansen, 2014). Maternal effect correlations (*r*_M_) between cold tolerance and heat tolerance (the only pair of traits for which this could be estimated) were also indistinguishable from zero and had similarly wide confidence intervals at both gradients (Table S1).

### Spatially variable selection on thermal tolerance and body size of *D. birchii* along elevation gradients

Linear selection differentials ranged from −0.46–0.63 (range |*S*| = 0.007 – 0.63; mean |*S*| = 0.172)(Table S2). Although none of the individual selection differentials was significantly different from zero after correction for multiple comparisons (Table S2), there was a strong and significant linear association between *S* and elevation for cold tolerance at Paluma (*R*^2^ = 0.685, *F*_1,8_ = 17.4, *P* = 0.003, *P_FDR_* = 0.019; Figure 2, Table 3). This association was negative, suggesting selection for increased cold tolerance at lower elevation sites (Figure 2). There was no significant linear association between *S* and elevation for any of the other traits at either gradient.

**Figure 2.**
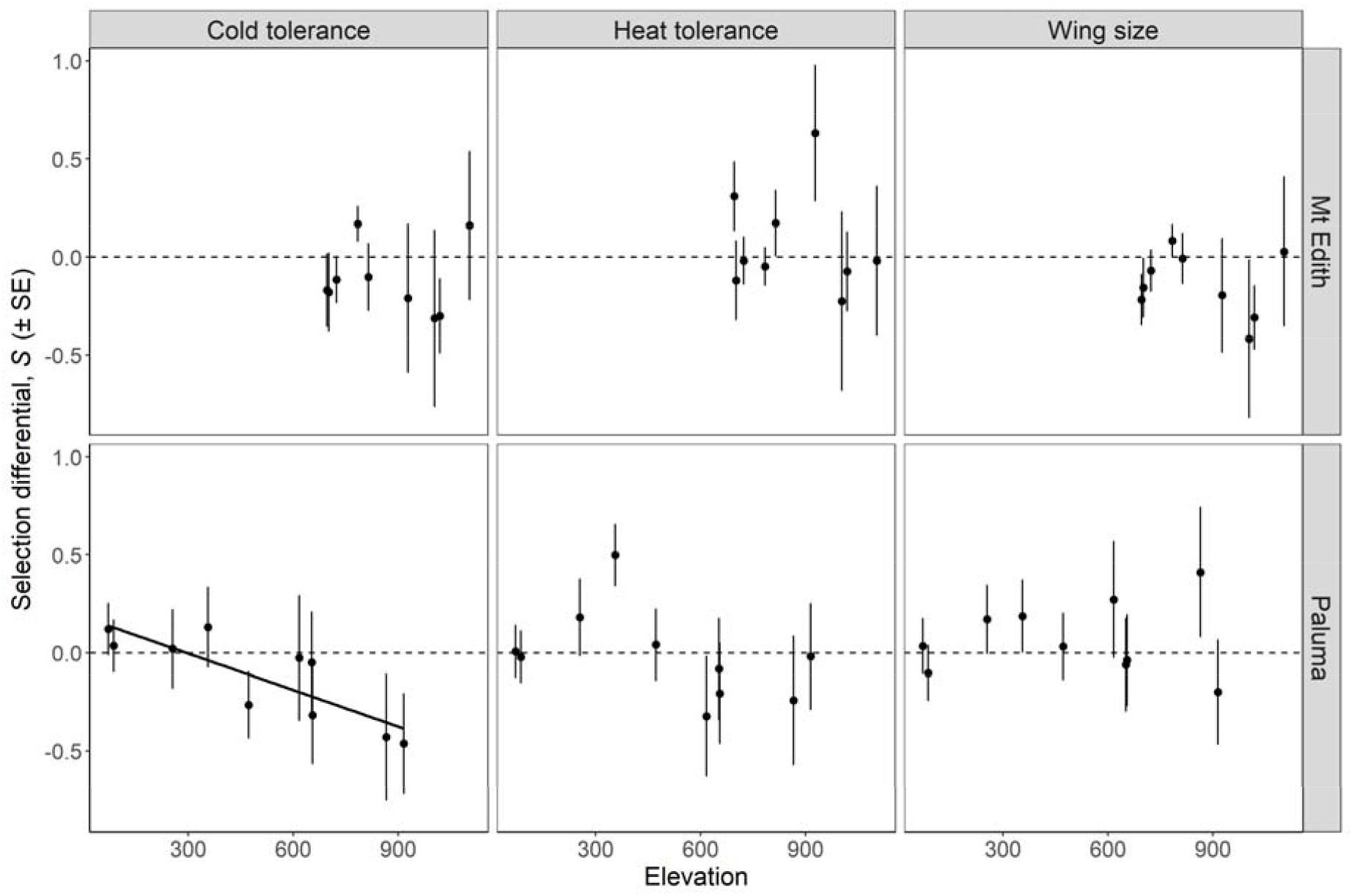
Linear selection differentials, *S*, for three traits of *Drosophila birchii* (cold tolerance, heat tolerance and wing size), estimated from the coefficients of regressions of standardised line means on relative fitness in caged transplant experiments for each trait at each site along two elevation gradients: Mt Edith (top row) and Paluma (bottom row). Error bars are standard errors of the regression coefficients at each site. The dashed line shows selection differential = 0 (no predicted change due to selection), therefore points above the line indicate positive directional selection and points below the line indicate negative directional selection. We tested for a linear relationship of selection differential with elevation along each gradient for each trait (see Table S4). This was only significant for cold tolerance at Paluma, where selection differentials declined with elevation, implying selection for lower cold tolerance at higher elevations (*R*^2^ = 0.685, *F*_1,8_ = 17.4, *P* = 0.003, *P_FDR_* = 0.019; Table S4).

**Table 3.** Results of regression models testing for a linear relationship between elevation and estimated selection differentials (*S*)(from Table S2) for each trait (cold tolerance, heat tolerance and wing size) at each gradient. *N* is the number of sites along each gradient, *β* is the slope of the relationship of *S* with elevation and SE is the standard error of this relationship, estimated from the linear model. Also shown is the result of an F-test (*F*_1,n-2_) and its associated p-value (*P*). *P_FDR_* is the p-value following correction for multiple tests using the False Discovery Rate. Traits where a significant association with elevation was seen after FDR correction (*P_FDR_* <0.05) are shown in bold italics. At Paluma, there was a strong negative linear association of *S* for cold tolerance with elevation; i.e. selection for reduced productivity following a cold shock at higher elevation (*R^2^* = 0.685; *F*_1,8_ = 17.36; *P* = 0.003; *P_FDR_* = 0.019).

| Gradient | Trait | $N$ | $R^2$ | $\beta$ | SE | $F_{(1,N-2)}$ | $P$ | $P_{\text{FDR}}$ |
| --- | --- | --- | --- | --- | --- | --- | --- | --- |
| Mt Edith | **Cold tolerance** | 9 | 0.000 | -4.264<br>$\times 10^{-7}$ | 4.317<br>$\times 10^{-4}$ | 0.000 | 0.999 | 0.999 |
| | **Heat tolerance** | 9 | 0.017 | -2.232<br>$\times 10^{-4}$ | 6.436<br>$\times 10^{-4}$ | 0.120 | 0.739 | 0.964 |
| | **Wing size** | 9 | 0.054 | -2.485<br>$\times 10^{-4}$ | 3.927<br>$\times 10^{-4}$ | 0.400 | 0.547 | 0.964 |
| Paluma | **Cold tolerance** | **10** | **0.685** | **-6.201**<br>**$\times 10^{-4}$** | **1.488**<br>**$\times 10^{-4}$** | **17.36** | **0.003** | **0.019** |
| | **Heat tolerance** | 10 | 0.214 | -3.625<br>$\times 10^{-4}$ | 2.457<br>$\times 10^{-4}$ | 2.176 | 0.178 | 0.535 |
| | **Wing size** | 10 | 0.008 | 5.686<br>$\times 10^{-5}$ | 2.207<br>$\times 10^{-4}$ | 0.066 | 0.803 | 0.964 |

### Change in heritability of wing size along elevation gradients, estimated from correlation of laboratory-reared females and their field-reared relatives

Gradient-wide wing size heritabilities estimated from lab field regressions at each transplant site were lower at Mt Edith (overall *H*^2^ = 0.05 (SE = 0.06); Table S3) but much higher at Paluma (overall *H*^2^ = 0.50 (SE = 0.06); Table S3), compared with those estimated from the laboratory quantitative genetic breeding design (Mt Edith *h*^2^ = 0.262, Paluma *h*^2^ = 0.292; Table S1). Individual site estimates exceeded lab estimates of *h*^2^ at most sites, although none of these estimates was significant after correction for multiple comparisons (Table S3). At both gradients, the highest field heritabilities were seen at the highest elevation site (Table S4). However, there was substantial variation in *H*^2^ within each gradient, even between adjacent sites, with field-estimated *H*^2^ ranging from 0.18–1 at Mt Edith and 0.07–1 at Paluma (Table S3). We did not detect a linear relationship between heritability and elevation at either gradient (Mt Edith: *R*^2^ = 0.383, *F*_1,5_ = 3.107, *P* = 0.138; Paluma: *R*^2^ = 0.019, *F*1,8 = 0.153, *P* = 0.706)(Table S4).

A regression of field *H*^2^ for wing size against the estimated selection differentials for this trait (*S*) at each gradient revealed a significant negative relationship between these parameters at Mt Edith (*R*^2^ = 0.771, *F*_1,5_ = 16.79, *P* = 0.009), but no significant relationship between them at Paluma: *R*^2^ = 0.042, *F*_1,8_ = 1.40, *P* = 0.271)(Figure 3; Table S5). Selection differentials for wing size at Mt Edith were all negative or very weakly positive (Table S2), therefore this result indicates more strongly negative selection (i.e. selection for smaller wing size) as *H*^2^ increases.

**Figure 3.**
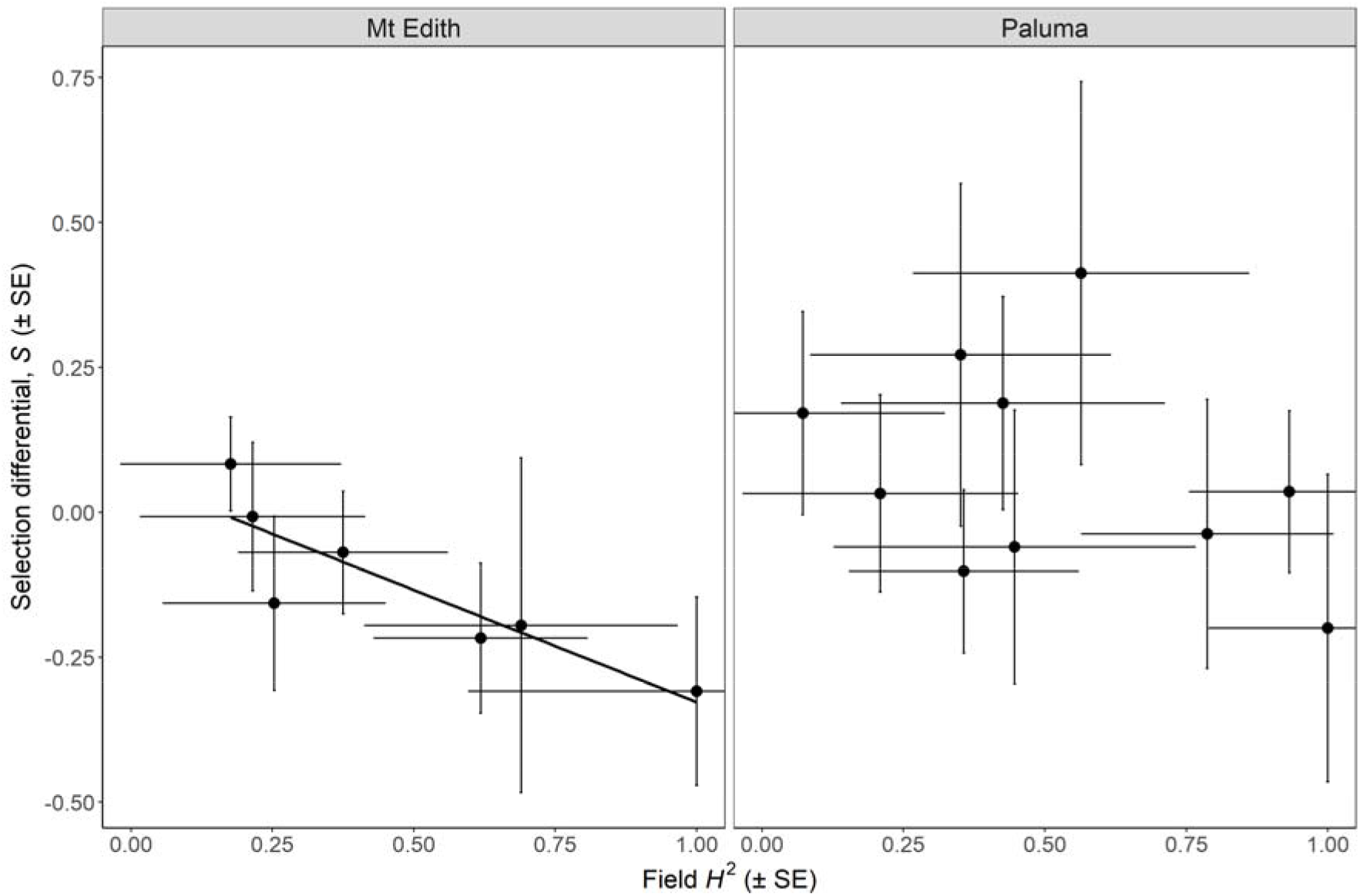
Plot of estimated selection differential (*S*) on wing size against field estimates of heritability (*H*^2^) of this trait at sites along two elevation gradients (Mt Edith and Paluma). Error bars are the standard errors of each of these estimates. There was a significant, negative relationship between *S* and *H*^2^ at Mt Edith (*R*^2^ = 0.771, *F*_1,5_ = 16.79, *P* = 0.009), indicating increasingly negative selection (i.e. selection for smaller wing size) where heritability is higher. The relationship was not significant at Paluma. (Table S5).

### Predicted response to selection on wing size along elevation gradients

The predicted response to selection on wing size (calculated using the breeder’s equation) at most sites along both gradients was the same or greater when using field *H*^2^ estimates compared with lab *h*^2^ estimates, albeit with wider errors (Figure 4). There was no significant linear relationship between elevation and predicted selection response *R* when using field *H*^2^ estimates (Mt Edith: *R*^2^ = 0.503, *F*_1,5_ = 5.064, *P* = 0.074; Paluma: *R*^2^ = 0.001, *F*_1,8_ = 0.006, *P* = 0.939; Table 4). However, there was a significant positive linear relationship between elevation and the magnitude of the predicted response to selection |*R*| at Paluma (*R*^2^ = 0.477, *F*_1,8_ = 7.284, *P* = 0.027). That is, a larger response to selection (ignoring direction) was predicted at higher elevation. While this relationship was not significant at Mt Edith, where fewer sites were available, the pattern was in the same direction and with a similar effect size (*R*^2^ = 0.528, *F*_1,5_ = 5.587, *P* = 0.064; Figure 4; Table 4).

**Figure 4.**
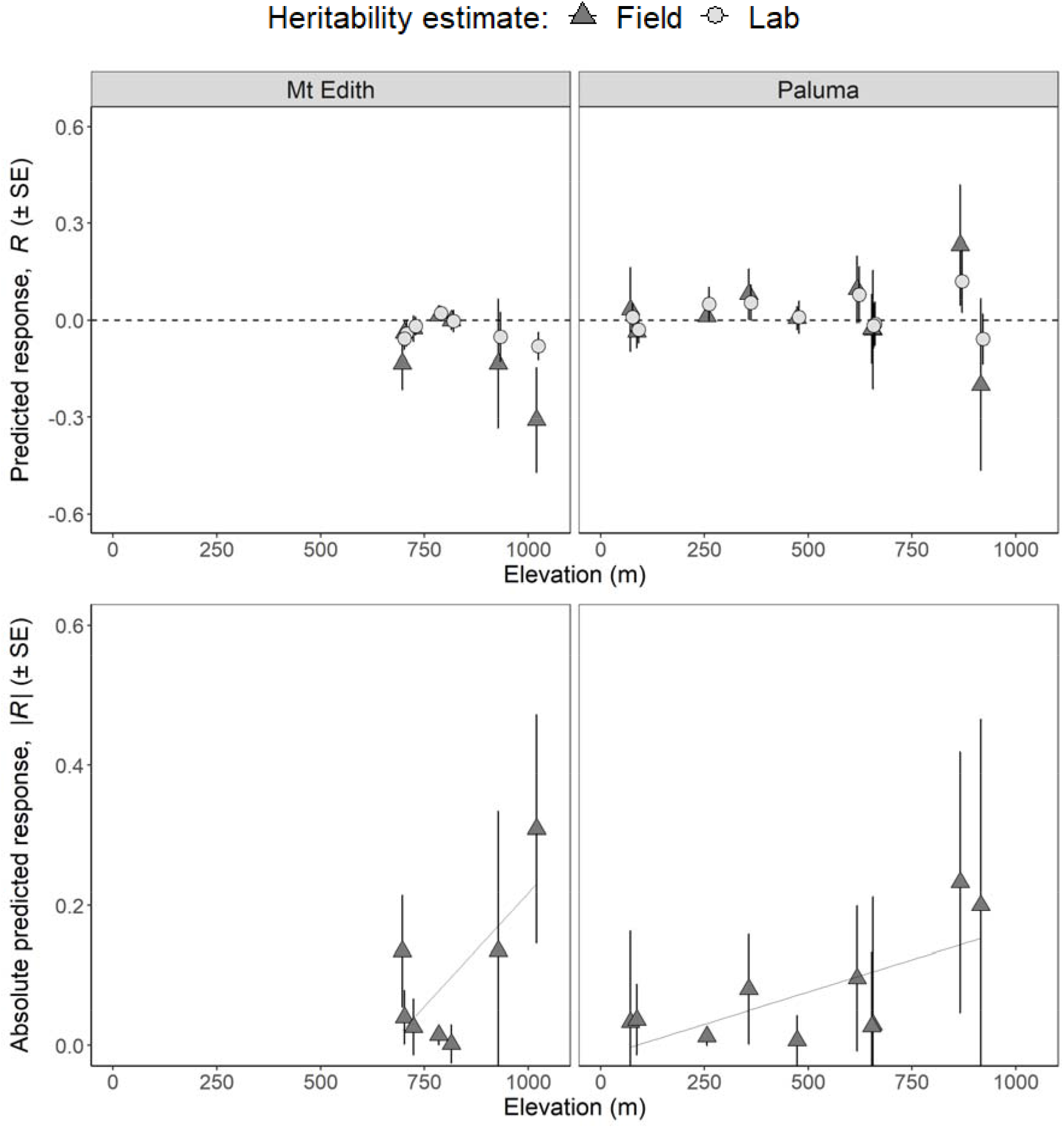
Predicted response to selection (in standard deviations (SDs) per generation) on wing size at sites along the two elevation gradients used in caged transplant experiments: Mt Edith (left) and Paluma (right). TOP: Predicted response to selection at each site was calculated using the breeders’ equation, as the product of the selection differential *S* as estimated from the covariance of wing size and fitness in field cages (see Figure 2; Table S3) and the trait heritability estimated from lab crosses (light grey circles; see Table S2) or parent-offspring correlations in field cages (dark grey triangles; see Table S5). BOTTOM: Absolute value of predicted selection response plotted against elevation for selection response estimates using field-based heritability estimates. There was a positive linear association of the absolute value of the selection response with elevation at both gradients, although this was significant only at Paluma (Mt Edith: *β*= 6.472 x 10^-4^, *F*_1,5_ = 5.587, *P* = 0.064; Paluma: β=1.842 x 10^-4^, *F*1,8 = 7.284, *P* = 0.027)(Table 4).

**Table 4.** Results of regression models testing for a linear relationship of elevation with (a) the predicted response to selection (*R)* on wing size; and (b) the absolute magnitude of the predicted response (i.e. regardless of direction; |*R*|) at each gradient. *R* was calculated using the breeder’s equation, as the product of *S* and *H*^2^ at each site along each gradient. *N* is the number of sites along each elevation gradient, *β* is the slope of the relationship of *R* or |*R*| with elevation and SE is the standard error of this relationship, estimated from the regression model. Also shown is the result of an F-test (*F*_1,n-2_)and its associated p-value (*P).* There was a positive relationship between the magnitude of the predicted selection response |*R*| and elevation at both gradients, which was significant at Paluma.

**(a) Regression of the predicted response to selection on wing size $R$ on elevation at each gradient**
| Gradient | $N$ | $R^2$ | $\beta$ | SE | $F_{(1,N-2)}$ | $P$ |
| --- | --- | --- | --- | --- | --- | --- |
| Mt Edith | 7 | 0.503 | $-6.553 \times 10^{-4}$ | $2.912 \times 10^{-4}$ | 5.064 | 0.074 |
| Paluma | 10 | 0.001 | $-1.039 \times 10^{-5}$ | $1.314 \times 10^{-4}$ | 0.006 | 0.939 |

**(b) Regression of the magnitude of the predicted response to selection on wing size $|R|$ on elevation at each gradient**
| Gradient | $N$ | $R^2$ | $\beta$ | SE | $F_{(1,N-2)}$ | $P$ |
| --- | --- | --- | --- | --- | --- | --- |
| Mt Edith | 7 | 0.528 | $6.472 \times 10^{-4}$ | $2.738 \times 10^{-4}$ | 5.587 | 0.064 |
| Paluma | 10 | **0.477** | **$1.842 \times 10^{-4}$** | **$6.824 \times 10^{-5}$** | **7.284** | **0.027** |

## Discussion

Understanding how the environment affects both selection on and heritability of traits in natural populations is critical for predicting the rate and direction of evolutionary responses to rapid environmental change (Husby et al., 2011, Hayward et al., 2018, Ramakers et al., 2018). By transplanting families of *Drosophila birchii* with known phenotypes in three traits to sites across two elevational gradients and measuring their fitness in the field, we were able to estimate selection at each site and test how it varies with elevation. For one of the traits (wing size), we estimated site-specific heritability by regressing wing size of offspring emerging from cages at field sites on that of their laboratory-reared relatives. We found variation in directional selection on cold tolerance at one gradient that should act to drive divergence of this trait with elevation. We also found that the size of the predicted response to selection on wing size increased at higher elevations. Together, these results suggest that evolutionary potential (both in terms of the strength of selection, and the likely response) can vary substantially along thermal gradients. This has important implications for the effects of environmental change in time and space on the capacity of populations of different species to adapt, and therefore to persist in ecological communities at given locations across their geographical ranges. Interestingly however, we did not observe divergence of trait means for any of the traits assayed between populations of *D. birchii* sourced from elevational extremes, suggesting that there are other factors constraining responses to selection in the field.

### Environmental variation in selection on cold tolerance, but not heat tolerance

Although overall selection differentials were similar for heat and cold tolerance of *D. birchii* within each gradient (Table S2), only cold tolerance showed spatial variation in selection that would drive divergence of this trait across the species’ elevational range (Figure 2; Table 3). Our estimates of linear selection differentials suggest that selection should drive *decreased* cold tolerance of *D. birchii* with increasing elevation, at least at the Paluma gradient. Selection for decreased cold tolerance at the cooler end of a species’ range is surprising, but perhaps suggests trade-offs with (unmeasured) traits that have a greater effect on fitness at high elevations. The method used to assay colder tolerance here was reproductive output immediately following a cold shock, so an alternative explanation for this unexpected result is that delaying reproduction after an acute stress can be adaptive in some cases (e.g. Wessels et al., 2011). Divergence of cold tolerance across elevation has previously been observed in *D. birchii* (Bridle et al., 2009); however, in that case, the measure was time to recover from a cold shock, which was in the direction usually expected i.e. flies sourced from cooler, high elevation sites recovered faster.

Studies across many taxa have found that lower thermal limits show much more variability within and among species than upper thermal limits, suggesting that cold tolerance has the potential to evolve more rapidly than heat tolerance (Bennett et al., 2021, Araújo et al., 2013, Hoffmann et al., 2013, Kinzner et al., 2019). In our transplant experiment, selection for heat tolerance may also be masked by the absence of biotic interactions in cages. There is growing empirical evidence that biotic interactions among species are a more important determinant of fitness at the warm edge of species’ ranges (Louthan et al., 2015, Paquette & Hargreaves, 2021, Zvereva & Kozlov, 2022). Our own assays of the effect of biotic interactions within cages on fitness of *D. birchii* at different elevations (O’Brien et al., 2020), as well as interactions with parasitoids or pathogens (Jeffs et al., 2021), also show increases in intensity at lower elevations. The absence of biotic interactions in cages may smooth out the environmental gradient with respect to the relationship of heat tolerance with fitness (i.e., fitness differences with respect to heat tolerance may only become evident in the presence of biotic stress). By contrast, the cage environment may exacerbate fitness variation associated with cold tolerance because it reduces the opportunity for flies to behaviourally thermoregulate (e.g., by seeking out more hospitable microclimates), or to actively forage for mates or oviposition sites, all of which are behaviours that are highly sophisticated in *Drosophila* (e.g. Churchill et al., 2020, Churchill et al., 2021, Malek & Long, 2020). Such behaviours would reduce variation in the strength of selection across the gradient, therefore the spatial variation in selection on cold tolerance that we detect from our estimates of cage fitness may be less pronounced under natural conditions.

### Predicted response to selection on wing size at the cold end of the elevational range of *D. birchii*

We saw a positive linear relationship between elevation and the magnitude of the predicted response to selection on wing size in *D. birchii* at Paluma, and a similar (though non-significant) relationship at Mt Edith (Table 4). This again suggests that the potential for evolutionary change in this trait increases towards the cooler, high elevation margin of *D. birchii’s* distribution, at least when flies and larvae are confined to cages and where interactions with other species are minimised. However, although the magnitude of the predicted response to selection increased with elevation, there was variation in direction, even between nearby sites, meaning that divergence of this trait was not predicted. Such divergence was also not observed.

### Environmental covariance of selection and heritability

Environmental coupling between selection and heritability determines evolutionary responses to environmental change in many cases, although tests of this are rare (Husby et al., 2011, Ramakers et al., 2018, Wilson et al., 2006, Wood & Brodie III, 2016). It also remains difficult to predict the direction of any such covariance, because although the strength of selection should increase in more stressful environments (Endler, 1986), the effect of stress on additive genetic variance and heritability is highly variable (Charmantier & Garant, 2005, Hoffmann & Merilä, 1999). Empirical studies have shown both a positive (Husby et al., 2011) and negative (Wilson et al., 2006) covariance of heritability and selection, with opposite consequences for the expected rate of evolutionary change. In our transplant experiments, we saw significant covariance of heritability and selection on wing size (the only trait for which we had site-specific heritability estimates) across sites at one of the elevation gradients (Mt Edith; Figure 3). This should increase the rate of evolutionary change in this trait. However, as noted above, the direction of selection was highly variable even within elevations and we saw no evidence for divergence of this trait with elevation.

### Other constraints to adaptation in the field

Our selection estimates are based on fitness estimated over a single generation, from (virgin) mating to egg to adult, whereas responses to selection in natural populations are effectively averages across many generations, which include temporal fluctuations in the strength (and possibly direction) of selection (Benning et al., 2021). Temporal variation in the environment can constrain evolutionary responses to directional selection due to variation in selection strength on traits during periods where conditions are more benign (Hao et al., 2015). However, experiments have also found positive effects of such temporal fluctuations on adaptation where population size increases during benign periods (Holt et al., 2004, Schaum et al., 2018). In the tropical mountain habitats considered here, there is much greater temporal variation in temperature and humidity (and presumably in biotic interactions) at low elevations than at high elevations (Saxon et al., 2018). Therefore, temporal variation may also contribute to the observation that cold tolerance frequently evolves more readily than heat tolerance (Kinzner et al., 2019, Hoffmann et al., 2013), if directional selection at cooler, more constant, high elevation sites is more effective.

The extent to which spatial variation in selection can drive genetic divergence of traits along a gradient also depends on the strength of selection relative to gene flow (Lenormand, 2002, Bridle & Vines, 2007, Hendry et al., 2002). Gene flow at this scale (between sites 1–10 km apart) is likely to be high in *D. birchii,* given the low differentiation observed at microsatellite markers between populations separated by hundreds of kilometres (Schiffer et al., 2007). Spatial variation in selection at such a local scale may therefore be small relative to the effect of gene flow between sites, particularly where reductions of population density at high and low elevations mean that gene flow is likely to be asymmetrical and therefore have strong swamping effects on local allele frequencies (O’Brien et al., 2017, 2022).

### Conclusion

Using a large cage transplant experiment, we were able to assess how evolutionary potential of three key traits varies across thermal gradients in the tropical fly *Drosophila birchii*. We found evidence for divergent selection on cold tolerance, but not heat tolerance or wing size. Furthermore, for wing size, where we could estimate environmental variation in responses to selection at sites along these gradients, we found that predicted responses were greater towards the cooler, high elevation edge of the gradient. Together, these results imply that there is greater potential for adaptation to conditions at the cold than the warm margin of this species’ range, which is concerning in light of expected warming under climate change.

However, divergence of traits between populations from elevational extremes was not observed, even when predicted from selection estimates. Failure to reconcile patterns of selection over the short timespan of our cage transplant experiment with patterns of trait variation in natural populations highlights the need for longer-term observations of selection in the wild, including consideration of other components of environmental variation such as biotic interactions.

## Acknowledgements

Peter Alexander, Chris Clinton, Ciara Mann and Lara Meade assisted with the field transplant experiments. Rachel Taylor, Alice Greaves, Calum Pennington and Tim Perry assisted with assays of cold and heat tolerance. Gareth Hobbs, Alexander Kelly, Laura Brooks, Laura Perkins, Holly So, Ali Somerville and Maddie Marzola assisted with mounting and measuring wings. Thank you to Greg Walter for helpful feedback on this manuscript. This work was funded by Natural Environment Research Council grant no. NE/G007039/1 to JRB.

## Data accessibility statement

All data and code used in the analysis presented in this manuscript can be found at the following link: https://github.com/EleanorOBrien/Estimating-change-in-evolutionary-potential-along-elevation-gradients.git

