## Supplementary Information for "Variation in the strength of selection but no trait divergence between elevational extremes in a tropical rainforest *Drosophila*"

**Appendix S1: Line cross design**

We established fully reciprocal diallel crosses among lines from within each of the eight source populations from Mt Edith and Mt Paluma. This crossing design is described in detail in O'Brien et al. (2017) and is summarised briefly here. All crosses were between lines within a given source population. We paired flies from each of the five lines in every combination, including within-line crosses, giving 25 combinations per population. Each cross was set up by placing one virgin male and female in a vial with 10 ml standard *Drosophila* media (agar, sugar, inactivated yeast, nipagin and propionic acid) for 4 days, then discarding them and leaving their offspring to emerge. This laying period results in a low density of offspring in each vial, thereby minimising effects of larval competition on traits. The crosses were set up in two separate generations: the first were established after lines had been in the laboratory for three generations, and offspring of these crosses were used in the field cage transplant experiment (see below and O'Brien et al. (2017)) and in assays of body size. The second set of crosses was established after the lines used for the field transplants had been in laboratory culture for six generations, and offspring of these crosses were used in assays of cold tolerance and heat tolerance. We established two replicate crosses per line combination for the first set of crosses, and three replicates for the second set.

Offspring emerging from the crossing design were sexed on the day of emergence under light CO_2_ anaesthesia, and held in single sex vials (maximum of 10 flies per vial) until they were used in either caged transplant experiments (3–10 days) or trait assays (6–7 days).

**Appendix S2: Measurement of quantitative traits**

*Cold tolerance*

We assayed cold tolerance in flies emerging from the diallel crossing design, to estimate additive genetic variance in this trait and to use in selection analyses (see main text).

Cold tolerance was assayed on six-day-old, previously unmated, females as fecundity following a cold shock. We placed flies individually in 40 ml vials containing 10 ml standard *Drosophila* media, then packed the vials into an ice-filled tray and placed them in a 4**°**C refrigerated room for 2 hrs. Following cold shock, we allowed flies 1 hour to recover at 25**°**C, then presented each female with a male taken from a mass-bred stock population that had been created by mixing flies from each line. We left flies to mate and lay for 72 hrs, then discarded them and left their offspring to develop. We recorded the number of offspring produced by each female as the measure of fecundity following cold shock. We assayed up to three females per family from the diallel breeding design (Mt Edith: 165 families, 419 flies; Paluma: 174 families, 485 flies).

*Heat tolerance*

We assayed heat tolerance in seven-day-old virgin females emerging from the diallel crossing design, to estimate additive genetic variance in this trait and to use in selection analyses (see main text). Heat tolerance was assayed as the time taken for flies to lose the ability to remain standing when exposed to a static heat shock of 37**°**C. We followed procedures similar to those described in Mitchell and Hoffmann (2010): we placed flies in empty 7 mL glass bottles (2 flies per bottle), which were secured with watertight lids and submerged in a water bath fitted with a thermoregulator (Ratek) that enabled the water temperature to be finely controlled. We recorded the time at which flies were no longer able to stand to the nearest second. We assayed up to three females per family from the diallel breeding design (Mt Edith: 188 families, 411 flies; Paluma: 199 families, 449 flies).

*Wing size*

We used wing size as a proxy for body size, since it is generally highly correlated with other body measurements in *Drosophila* (Gilchrist et al., 2001). We measured wing size on three sets of *D. birchii* to address different objectives of our study: we assayed wing size in flies emerging from the same set of diallel crosses that provided the source flies for the caged transplant experiment to estimate additive genetic variance of this trait, and in analyses of selection (see main text); we also assayed wing size of (2) surviving female *D. birchii* and (3) their female offspring emerging from field cages in the transplant experiment, to estimate heritability of this trait at different sites along elevation gradients. The same protocol was used in all cases.

We measured the right wing of female flies that were at least two days old. Wing size is fixed in adult flies and so is not affected by age, but we aged them for two days to ensure their wings were completely unfurled. Assays of wing size were identical to those described in (Saxon et al., 2018). Briefly, the wing was removed, mounted on a glass slide with Aquatex mounting agent, and covered with a glass cover slip. Each wing was photographed at 20 X magnification using a digital camera (GXCAM-9) mounted on a Nikon SMZ800 microscope. We used the program *tpsDig2* (available at http://life.bio.sunysb.edu/morph/) to add landmarks to wing images at 10 positions corresponding to vein intersections or terminations (Figure S1), following the method of Griffiths et al. (2005). Wing size was measured as centroid size (in mm), which is the square root of the sum of the squared distances between each landmark and the mean position of all of the landmarks (the wing centroid). Centroid sizes were obtained using *MorphoJ* (Klingenberg, 2011). We excluded any damaged wings (i.e. those lacking any of the landmarks). We re-photographed and landmarked 10% of wings in the data set to obtain an estimate of error. Variation among repeat measurements of the same wings was < 1% of total variation.

We assayed wing size for an average of 8‒10 females per family from the diallel crosses (Mt Edith: 35 families, 345 flies; Paluma: 35 families, 283 flies). There were fewer families assayed for this trait because (1) only two replicates of each line cross were established in this round of crosses and (2) flies were preferentially used to establish the caged transplant experiment, which meant there were not sufficient flies remaining from all families to assay wing size. We also measured wing size of all of the females used to found the caged transplant experiment that were still surviving after five days in the field (Mt Edith: 362 flies; Paluma: 282 flies) and their female offspring emerging from the field cages (Mt Edith: 183 flies; Paluma: 174 flies).

**Appendix S3: Pedigree construction for estimating additive genetic effects in *MCMCglmm***

We used the pedigree structure from our breeding design to generate a pairwise relatedness matrix for each gradient, giving the relatedness coefficient between each pair of individuals. We made the following assumptions about the relatedness of individuals within isofemale lines: (1) the first generation of offspring in each line were full-siblings, i.e., offspring produced by a field-mated female were all sired by a single male. This assumption fits with our understanding of the biology of this species, where females lack spermathecae and so have limited capacity to store sperm, females resist remating within 24 hours (personal observation), and there is strong last male precedence in siring success when females are mated multiply (unpublished data), (2) the parents of the isofemale lines were no more related to one another than two individuals selected at random from the population, and (3) lines had been maintained at a mean population size of 200 individuals per generation for six generations. This gives an inbreeding coefficient (*F*) of 0.257 within each line at the time when the breeding design was established. Flies assayed for body size had only been maintained for three generations, meaning the inbreeding coefficient was marginally smaller when they were assayed (*F* = 0.252). However, this made no discernible difference to the estimate of additive genetic variance in this trait in univariate analyses, and for the multivariate models it was simpler to assume a constant value of *F*. We therefore assumed *F* = 0.257 within lines when constructing the pedigree for models including all three traits.

**Supplementary Tables**

**Table S1**. (a) Heritabilities (on the diagonal, in bold) of traits and genetic correlations (below the diagonal, in italics) among traits and (b) Maternal effect variance as a proportion of total phenotypic variance (on the diagonal, in bold) of traits and maternal effect correlations (below the diagonal, in italics) among traits for *D. birchii* from Mt Edith and Paluma, estimated from MCMCglmm models, with upper and lower 95% Highest Posterior Density (HPD) intervals for each estimate. Note that maternal effect correlations were not estimated for wing size because this trait was measured on offspring from different mothers to cold and heat tolerance.

| (a) **Heritabilities and genetic correlations** | | | | | | | | |
| --- | --- | --- | --- | --- | --- | --- | --- | --- |
|  | **Mt Edith** | | |  |  | **Paluma** | | |
| **Trait** | **Cold tolerance** | **Heat tolerance** | **Wing**  **size** |  | **Trait** | **Cold tolerance** | **Heat tolerance** | **Wing**  **size** |
| **Cold tolerance** | **0.199**  **(0.06, 0.35)** |  |  |  | **Cold tolerance** | **0.167 (0.06, 0.29)** |  |  |
| **Heat tolerance** | *0.086*  *(-0.43, 0.54* | **0.117 (0.05, 0.19)** |  |  | **Heat tolerance** | *0.036*  *(-0.42, 0.49)* | **0.168 (0.08, 0.26)** |  |
| **Wing**  **size** | *0.002*  *(-0.60, 0.67)* | *-0.008*  *(-0.54, 0.54)* | **0.262 (0.06, 0.56)** |  | **Wing**  **size** | *0.066*  *(-0.52, 0.64)* | *-0.029*  *(-0.61, 0.52)* | **0.292 (0.09, 0.53)** |
| (b) **Maternal effect variance and correlations** | | | | | | | | |
|  | **Mt Edith** | | |  |  | **Paluma** | | |
| **Trait** | **Cold tolerance** | **Heat tolerance** | **Wing**  **size** |  | **Trait** | **Cold tolerance** | **Heat tolerance** | **Wing**  **size** |
| **Cold tolerance** | **0.167 (0.07, 0.26)** |  |  |  | **Cold tolerance** | **0.150 (0.07, 0.24)** |  |  |
| **Heat tolerance** | *0.115*  *(-0.29, 0.54)* | **0.098 (0.05, 0.15)** |  |  | **Heat tolerance** | *-0.01*  *(-0.44, 0.37)* | **0.090 (0.05, 0.15)** |  |
| **Wing**  **size** | - | - | **0.223 (0.06, 0.41)** |  | **Wing**  **size** | - | - | **0.170 (0.05, 0.29)** |

**Table S2**. Selection differentials (*S*) and their standard error (SE) for each trait at each site along the Mt Edith and Paluma gradients, calculated as the slope of a regression of standardised trait mean against relative fitness at each site. Fitness was the productivity of flies transplanted in cages at each site. *N*_cage_ is the number of cages for which we had a measure of fitness, and *N*_line_ is the number of maternal isofemale lines for which we estimated trait means at each site. *P*-values for each regression, and *P*-values corrected for multiple tests using the False Discovery Rate (*P_FDR_*) of (Benjamini & Hochberg, 1995) are also shown. None of the individual values of *S* was significantly different from zero after correcting for multiple tests. Also shown is the overall selection differential estimated for each gradient for each trait, which was also not significant for any of the traits at either gradient.

| **Trait** | **Mt Edith** | | | | | | | **Paluma** | | | | | | |
| --- | --- | --- | --- | --- | --- | --- | --- | --- | --- | --- | --- | --- | --- | --- |
|  | **Elevation** | ***N*_cage_** | ***N*_line_** | ***S*** | **SE** | ***P*** | ***P*_FDR_** | **Elevation** | ***N*_cage_** | ***N*_line_** | ***S*** | **SE** | ***P*** | ***P*_FDR_** |
| **Cold tolerance** | 697 | 20 | 5 | -0.17 | 0.19 | 0.37 | 0.72 | 72 | 32 | 14 | 0.12 | 0.13 | 0.360 | 0.717 |
|  | 702 | 19 | 5 | -0.18 | 0.20 | 0.38 | 0.72 | 87 | 31 | 14 | 0.04 | 0.13 | 0.790 | 0.936 |
|  | 724 | 19 | 6 | -0.11 | 0.12 | 0.34 | 0.72 | 256 | 17 | 6 | 0.02 | 0.20 | 0.921 | 0.936 |
|  | 785 | 20 | 5 | 0.17 | 0.09 | 0.08 | 0.66 | 357 | 16 | 7 | 0.13 | 0.20 | 0.529 | 0.794 |
|  | 815 | 19 | 5 | -0.10 | 0.17 | 0.55 | 0.79 | 473 | 17 | 6 | -0.26 | 0.17 | 0.140 | 0.664 |
|  | 928 | 17 | 5 | -0.21 | 0.38 | 0.59 | 0.79 | 618 | 15 | 5 | -0.03 | 0.32 | 0.936 | 0.936 |
|  | 1005 | 12 | 5 | -0.31 | 0.45 | 0.50 | 0.79 | 653 | 16 | 6 | -0.05 | 0.26 | 0.855 | 0.936 |
|  | 1020 | 18 | 5 | -0.30 | 0.19 | 0.13 | 0.66 | 656 | 17 | 7 | -0.32 | 0.25 | 0.218 | 0.689 |
|  | 1105 | 20 | 5 | 0.16 | 0.38 | 0.68 | 0.86 | 866 | 27 | 13 | -0.43 | 0.32 | 0.193 | 0.689 |
|  |  |  |  |  |  |  |  | 916 | 27 | 13 | -0.46 | 0.25 | 0.081 | 0.664 |
|  | **Overall** | **164** | **6** | **-0.08** | **0.06** | **0.200** | **0.400** | **Overall** | **215** | **14** | **-0.01** | **0.06** | **0.860** | **0.860** |
| **Heat tolerance** | 697 | 20 | 5 | 0.31 | 0.18 | 0.095 | 0.600 | 72 | 32 | 14 | 0.01 | 0.13 | 0.956 | 0.960 |
|  | 702 | 19 | 5 | -0.12 | 0.20 | 0.557 | 0.960 | 87 | 31 | 14 | -0.02 | 0.13 | 0.878 | 0.960 |
|  | 724 | 19 | 6 | -0.02 | 0.12 | 0.879 | 0.960 | 256 | 17 | 6 | 0.18 | 0.19 | 0.365 | 0.960 |
|  | 785 | 20 | 5 | -0.05 | 0.10 | 0.624 | 0.960 | 357 | 16 | 7 | 0.50 | 0.16 | 0.007 | 0.126 |
|  | 815 | 19 | 5 | 0.17 | 0.17 | 0.314 | 0.960 | 473 | 17 | 6 | 0.04 | 0.18 | 0.822 | 0.960 |
|  | 928 | 17 | 5 | 0.63 | 0.34 | 0.087 | 0.600 | 618 | 15 | 5 | -0.32 | 0.31 | 0.311 | 0.960 |
|  | 1005 | 12 | 5 | -0.23 | 0.45 | 0.630 | 0.960 | 653 | 16 | 6 | -0.08 | 0.26 | 0.756 | 0.960 |
|  | 1020 | 18 | 5 | -0.07 | 0.20 | 0.719 | 0.960 | 656 | 17 | 7 | -0.21 | 0.25 | 0.429 | 0.960 |
|  | 1105 | 20 | 5 | -0.02 | 0.38 | 0.960 | 0.960 | 866 | 27 | 13 | -0.24 | 0.33 | 0.466 | 0.960 |
|  |  |  |  |  |  |  |  | 916 | 27 | 13 | -0.02 | 0.27 | 0.948 | 0.960 |
|  | **Overall** | **164** | **6** | **0.10** | **0.06** | **0.126** | **0.378** | **Overall** | **215** | **14** | **0.03** | **0.06** | **0.642** | **0.770** |
| **Wing size** | 697 | 29 | 9 | -0.22 | 0.13 | 0.104 | 0.756 | 72 | 29 | 13 | 0.04 | 0.14 | 0.802 | 0.954 |
|  | 702 | 31 | 11 | -0.16 | 0.15 | 0.305 | 0.756 | 87 | 29 | 13 | -0.10 | 0.14 | 0.476 | 0.756 |
|  | 724 | 32 | 12 | -0.07 | 0.11 | 0.517 | 0.756 | 256 | 19 | 7 | 0.17 | 0.18 | 0.343 | 0.756 |
|  | 785 | 30 | 9 | 0.08 | 0.08 | 0.312 | 0.756 | 357 | 18 | 8 | 0.19 | 0.18 | 0.322 | 0.756 |
|  | 815 | 28 | 9 | -0.01 | 0.13 | 0.954 | 0.954 | 473 | 19 | 7 | 0.03 | 0.17 | 0.850 | 0.954 |
|  | 928 | 23 | 10 | -0.20 | 0.29 | 0.507 | 0.756 | 618 | 16 | 6 | 0.27 | 0.30 | 0.373 | 0.756 |
|  | 1005 | 13 | 6 | -0.42 | 0.40 | 0.322 | 0.756 | 653 | 18 | 7 | -0.06 | 0.24 | 0.803 | 0.954 |
|  | 1020 | 26 | 9 | -0.31 | 0.16 | 0.070 | 0.756 | 656 | 19 | 8 | -0.04 | 0.23 | 0.875 | 0.954 |
|  | 1105 | 20 | 5 | 0.03 | 0.38 | 0.942 | 0.954 | 866 | 26 | 13 | 0.41 | 0.33 | 0.224 | 0.756 |
|  |  |  |  |  |  |  |  | 916 | 26 | 13 | -0.20 | 0.27 | 0.459 | 0.756 |
|  | **Overall** | **232** | **12** | **-0.09** | **0.05** | **0.072** | **0.378** | **Overall** | **219** | **13** | **0.05** | **0.06** | **0.457** | **0.686** |

**Table S3.** Wing size heritability (*H*^2^) at each transplant site along the Mt Edith and Paluma elevation gradients, estimated as twice the slope of a regression of standardised body size of female offspring emerging from cages against standardised body size of laboratory-reared females from the same maternal isofemale line. Standard error (SE) is the SE of the fitted regression line. *N_cages_* is the number of cages at each site included in this analysis (i.e. cages where wing size measurements of both sets of flies used in the regression analysis were available). Heritability was not estimated for two sites at Mt Edith because too few cages had offspring emerging. Also shown are *P*-values of each regression and *P*-values corrected for multiple tests using the False Discovery Rate (*P_FDR_*) method of (Benjamini & Hochberg, 1995). Notionally significant (*P* < 0.05) *H*^2^ estimates are shown in italics. None of the individual site *H*^2^ estimates was significant after adjusting for multiple comparisons (*P_HDR_* > 0.05 in all cases). Also shown is the overall estimate of *H*^2^ of body size at each gradient, from regressions across all cages, with transplant site included as a random factor. This was highly significant at Paluma (*H*^2^ = 0.50, SE = 0.06, *P* = 2.97 x 10^-5^). Because true values of *H*^2^ will lie between 0 and 1, we estimated *H*^2^ = 1 when the estimate from the regression exceeded 1. These values are marked with a *.

| **Mt Edith** | | | | | | **Paluma** | | | | | |
| --- | --- | --- | --- | --- | --- | --- | --- | --- | --- | --- | --- |
| **Elevation (m)** | ***N_cages_*** | ***H*^2^** | ***SE*** | ***P*** | ***P_FDR_*** | **Elevation (m)** | ***N_cages_*** | ***H*^2^** | ***SE*** | ***P*** | ***P_FDR_*** |
| 697 | 24 | 0.62 | 0.19 | 0.11 | 0.46 | *72* | *12* | *0.93* | *0.18* | *0.01* | *0.07* |
| 702 | 26 | 0.25 | 0.20 | 0.52 | 0.65 | 87 | 17 | 0.36 | 0.20 | 0.39 | 0.65 |
| 724 | 13 | 0.38 | 0.18 | 0.32 | 0.56 | 256 | 16 | 0.07 | 0.25 | 0.89 | 0.89 |
| 785 | 10 | 0.18 | 0.19 | 0.65 | 0.65 | 357 | 16 | 0.43 | 0.29 | 0.47 | 0.65 |
| 815 | 11 | 0.22 | 0.20 | 0.59 | 0.65 | 473 | 17 | 0.21 | 0.24 | 0.67 | 0.75 |
| 928 | 26 | 0.69 | 0.28 | 0.24 | 0.56 | 618 | 13 | 0.35 | 0.26 | 0.52 | 0.65 |
| 1020 | 17 | 1.00* | 0.40 | 0.13 | 0.46 | 653 | 24 | 0.45 | 0.32 | 0.50 | 0.65 |
|  |  |  |  |  |  | 656 | 29 | 0.79 | 0.22 | 0.10 | 0.32 |
|  |  |  |  |  |  | 866 | 27 | 0.56 | 0.30 | 0.37 | 0.65 |
|  |  |  |  |  |  | *916* | *28* | *1.00** | *0.21* | *0.01* | *0.07* |
| **Overall** | **127** | **0.05** | **0.06** | **0.68** |  | ***Overall*** | ***199*** | ***0.50*** | ***0.06*** | ***2.97x10^-5^*** |  |

**Table S4.** Results of regression models testing for a linear relationship between elevation and field estimates of wing size *H*^2^ (from Table S3) at each gradient. *N* is the number of sites along each elevation gradient, *β* is the slope of the relationship of *H*^2^ with elevation and SE is the standard error of this relationship, estimated from the regression model. Also shown is the result of an F-test (*F*_1,n-2_) and its associated p-value (*P*). We did not find a significant linear association of *H*^2^ with elevation at either gradient.

| **Gradient** | ***N*** | ***R*^2^** | ***β*** | **SE** | ***F*_(1,_*_N_*_-2)_** | ***P*** |
| --- | --- | --- | --- | --- | --- | --- |
| **Mt Edith** | 7 | 0.383 | 1.582 x 10^-3^ | 8.977 x 10^-4^ | 3.107 | 0.138 |
| **Paluma** | 10 | 0.019 | 1.416 x 10^-4^ | 3.616 x 10^-4^ | 0.153 | 0.706 |

**Table S5.** Results of regression models testing for a linear relationship between the selection differential for wing size (S; from Table S2) and field estimates of body size *H*^2^ (from Table S3) at each gradient. *N* is the number of sites along each elevation gradient, *β* is the slope of the relationship of *S* with *H*^2^ and SE is the standard error of this relationship, estimated from the regression model. Also shown is the result of an F-test (*F*_1,n-2_) and its associated p-value (*P*). These data are plotted in Figure S3.

| **Gradient** | ***N*** | ***R*^2^** | ***β*** | **SE** | ***F*_(1,_*_N_*_-2)_** | ***P*** |
| --- | --- | --- | --- | --- | --- | --- |
| ***Mt Edith*** | ***7*** | ***0.771*** | ***-0.387*** | ***0.094*** | ***16.788*** | ***0.009*** |
| **Paluma** | 10 | 0.042 | -0.236 | 0.200 | 1.397 | 0.271 |

**Supplementary Figures**

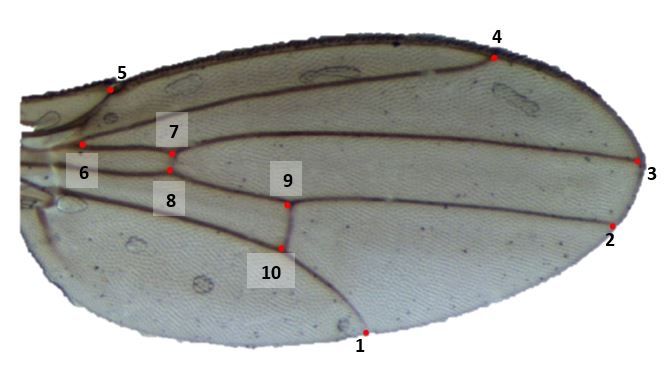

**Figure S1.** Position of the 10 landmarks used to calculate wing centroid sizes in *Drosophila birchii*.

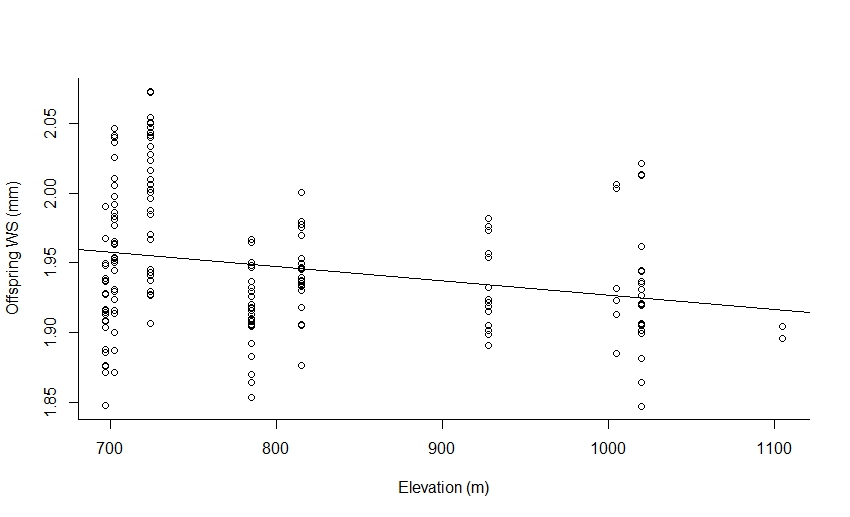

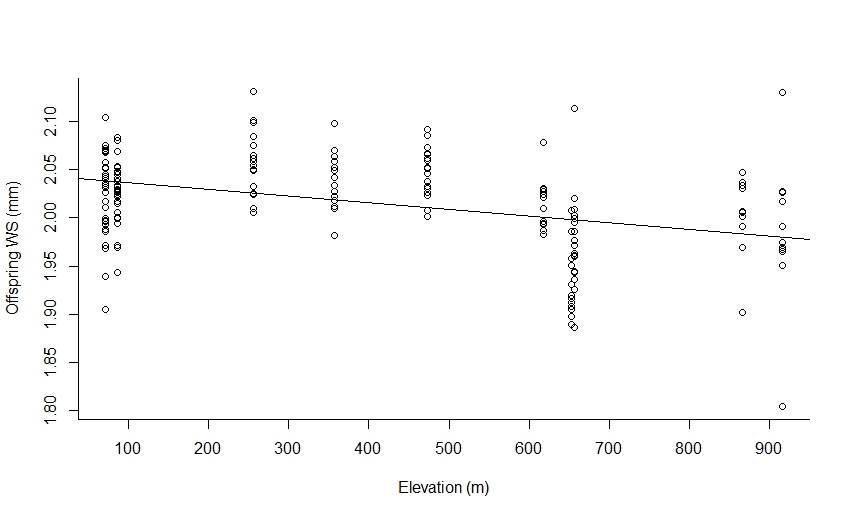

*R*^2^ = 0.142, *F*_1,172_ = 29.68, *P* <0.001

**Paluma**

*R*^2^ = 0.056, *F*_1,181_ = 11.76, *P* < 0.001

**Mt Edith**

**Figure S2.** Mean wing size (WS) of *D. birchii* female offspring emerging from cages at each transplant site along the Mt Edith (left) and Paluma (right) gradients. Points are cage means. We measured wing size of offspring emerging from 162 cages at Mt Edith and 174 cages at Paluma (see Table S5 for number of cages at each transplant site from which offspring were measured). The line on each plot is the fitted line from a regression of mean offspring wing size on elevation at each gradient. At both gradients, there was a significant negative relationship between elevation and the mean wing size of offspring emerging from cages.
